# High-efficiency digitally scanned light-sheet fluorescence lifetime microscopy (DSLM-FLIM)

**DOI:** 10.1101/2023.06.02.543377

**Authors:** Kyle J. Nutt, Daniel Olesker, Ewan McGhee, Graham Hungerford, Christopher G. Leburn, Jonathan Taylor

**Affiliations:** School of Physics and Astronomy, University of Glasgow, Glasgow G12 8QQ, UK; Cancer Research UK Beatson Institute, Garscube Estate, Glasgow G61 1BD, UK; HORIBA Jobin Yvon IBH Ltd, Glasgow G3 8HB, UK; Chromacity, Edinburgh EH14 4AP, UK

## Abstract

Recently-developed kilopixel single-photon avalanche diode (SPAD) arrays with in-pixel timing hold great promise for fluorescence lifetime imaging microscopy of dynamic samples, thanks to their widefield single-photon time-of-arrival imaging capabilities. However digitally-scanned light-sheet microscope (DSLM) and two-photon microscope systems present significant technical barriers which have to date prevented full and efficient use of the capabilities of SPAD arrays. Because the 12.4 kHz frame-rate of our array camera is faster than achievable DSLM scan rates, most pixels would be sitting idle most of the time. We present a new optical design based around astigmatic imaging optics, enabling rapid and efficient acquisition of fluorescence lifetime imaging data. We demonstrate our system with both one- and two-photon excitation sources, validate performance with lifetime reference beads, and demonstrate separation of similar fluorescence emission spectra in biological samples *via* lifetime contrast.

---

Fluorescence lifetime imaging microscopy (1) is a powerful tool for probing the chemical and biophysical microenvironment of biological cells. Chemical properties such as pH (2) and ion concentration (3) influence the excited state lifetime of a fluorescent probe, and molecular proximity, tension and conformational changes can be probed sensitively *via* changes in the fluorescence lifetime induced by Förster Resonance Energy Transfer (FRET) (4, 5). A powerful and potentially very sensitive method for imaging fluorescence lifetime is time-correlated single-photon counting (TCSPC) using a single-photon avalanche diode (SPAD) array: an array of avalanche photodiodes in the style of a conventional pixelated camera (6, 7). Each SPAD is associated with a dedicated time to digital converter (TDC), enabling each pixel to make picosecond-resolution measurements of the arrival time of individual emitted photons (we denote this timebase as *τ*). In the SPAD array architecture of the Horiba Flimera TCSPC camera (8), each pixel operates with effective exposure-windows of approximately 80 µs (we denote this timebase as *T*), within which each pixel is capable of registering a single time-tagged photon arrival. Although this can be likened to a conventional camera sensor capturing binary images at a frame-rate of approximately 12 kHz, the key feature of a SPAD camera is that, in addition to the *xy* pixel coordinates of each detected photon, it also records the precise time-of-arrival of each photon with a time-resolution of approximately 36 picoseconds, providing fluorescence lifetime information for each pixel in the image.

The massive parallelism of this type of SPAD array is ideal for increasing FLIM acquisition rates (9, 10), opening up realistic prospects for three-dimensional and time-resolved FLIM imaging in dynamic living samples, with higher photon efficiencies than gated CCD architectures (6). However, many FLIM experiments are based around point-or line-scanned excitation (confocal or two-photon microscopy with a pulsed laser source and a single SPAD detector). Indeed, in a two-photon scenario this is near-essential to ensure efficient two-photon excitation by concentrating the laser power into a more compact region in time and space, compared to wide-field excitation. (11). This introduces a fundamental tension between the specifications of SPAD array cameras and their application to time-resolved scan-based FLIM imaging of dynamic samples. Each individual detector in a SPAD array is read out with an effective frame-rate of tens of thousands of single-photon frames per second, and so any laser-scanned microscope (maximum scan rates typically a few kHz) will inevitably not make the most efficient use of the SPAD array. Only a small proportion of pixels will be illuminated, with the majority “idling ” most of the time.

A pertinent example of where this trade-off would be encountered is in digitally-scanned light-sheet imaging. Here, a pencil beam is scanned rapidly in the focal plane of the detection objective, forming an illumination profile that averages over the course of an exposure-window to a thin light-sheet. However, during the course of the exposure-window, the pencil beam is positioned in a sample region conjugate to each row of pixels for only a small fraction of the overall acquisition time. In conventional intensity imaging this does not matter, since each pixel typically integrates over exposures lasting milliseconds or seconds, with no restrictions on the temporal distribution of individual photon arrival times. However, the greatly increased temporal resolution offered by the SPAD array architecture means that each avalanche photodiode would be sitting idle for the majority of the overall acquisition, representing an inefficient use of the detector.

The tension between the efficient use of a widefield SPAD camera, taking advantage of its ability to work at rates oftens of thousands of detected photons per second, and the need for spatially-concentrated instantaneous laser excitation for efficient two-photon microscopy, has until now restricted the usefulness of SPAD cameras for two-photon and DSLM-based FLIM (12).

We have developed a new optical design that overcomes this barrier to enable efficient two-photon DSLM-FLIM. Our astigmatic optical design ensures every pixel in the SPAD array has the potential to receive photons at all times throughout the laser scan process, hence exploiting the full parallel bandwidth of the SPAD array and overcoming the limitations discussed above. Figure 1(a,c) illustrates the concept and optical schematic for our system. A standard DSLM light sheet is formed by scanning a laser beam laterally (in *y*) using a galvanometric mirror (GM) conjugate to the exit pupil of the launch objective, to launch a scanned pencil beam into the sample. The optical design for the imaging arm is image-forming in the *x* dimension (image sensor conjugate to the objective focal plane; CL2 *f* = 150 mm) but distributes the instantaneous line-excitation across the image sensor in the *y* dimension (image sensor conjugate to the objective pupil plane; demagnifying 4*f* relay CL1 and CL3 *f* = 150 and 50 mm).

**Fig. 1.**
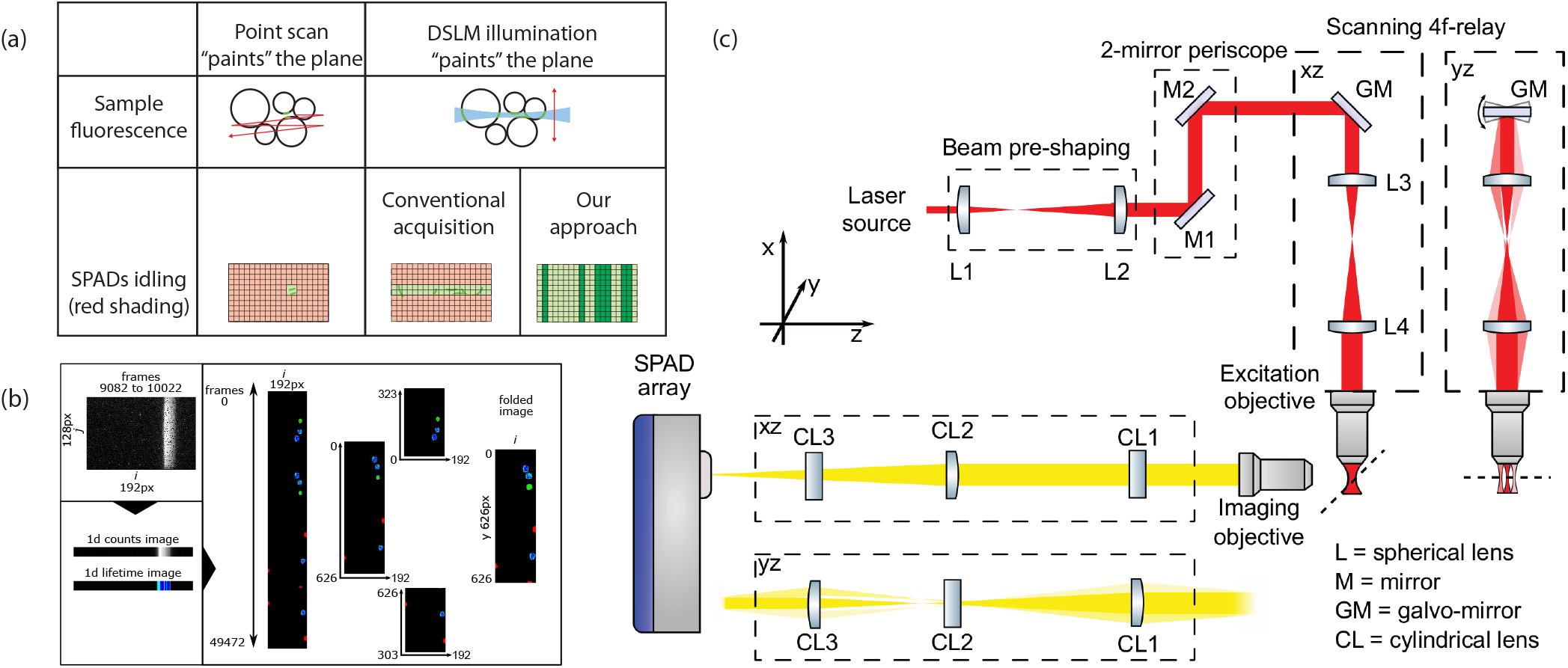
(a) **Concept**: point-scanned or conventional DSLM illumination would result in SPAD camera pixels idling most of the time, whereas our astigmatic imaging will ensure the SPAD camera is evenly illuminated throughout the DSLM scan. (b) **Processing pipeline from raw data gathered on the detector array to final images**. Batches of *n* raw binary frames are combined and summed vertically to yield an *xτ* time-of-arrival histogram for a single row of pixels in the final image. The final *xyτ* lifetime image is formed row-by-row in this manner. Mirror duplication about a horizontal axis (a consequence of the saw-tooth scan profile) is eliminated by folding, as described in the main text. (c) **Optical schematic**: a conventional DSLM microscope configuration excites fluorescence, which is imaged using our specialised astigmatic imaging configuration tailored to the SPAD sensor dimensions

We do not require to form an image on the sensor in the *y* dimension because the frame number (timebase *T*) for each photon event is known, and hence the instantaneous *y* position of the excitation beam in the sample can be easily inferred. This astigmatic imaging approach introduces two key benefits into our microscope. Firstly, unlike in DSLM where the fluorescence excited by the pencil beam at any one scan position is imaged to a narrow row of pixels, our system spreads the excited fluorescence across the entire height of the SPAD array detector. For any instantaneous position of the pencil beam the entire sensor could be actively detecting photons. This leads to a dramatically more efficient use of the detector than conventional DSLM. The increased efficiency effectively boosts the maximum instantaneous usable photon flux by up to two orders of magnitude, consequently achieving a corresponding reduction in the image acquisition time. Secondly, the field of view that can be imaged is extended in the vertical direction. Because the image is not focused along the vertical axis, our vertical field of view is not restricted by the extent or pixel count of the SPAD camera, and is only limited by the field of view of the objective lenses.

The process of reconstructing lifetime images from our astigmatic imaging system is outlined in Figure 1(b). Each raw frame records a series of single-photon time-of-arrival events. These can be considered as picosecond-resolved (and spatially resolved) delay measurements *τ*_*ijT*_ between excitation laser pulse generation and fluorescent photon emission (i.e. single-photon fluorescence lifetime). Alternatively these can be represented as a sparse event matrix *E*_*ijτT*_, where *τ* is the time-of-arrival (resolved into 4096 separate lifetime bins of width 36.6 ps), (*i, j*) denotes the pixel location on the sensor array, and *T* is the frame index (incremented at 12.4 kHz). In our optical setup the *i* coordinate of the event in the raw frame directly encodes the *x* coordinate in the output image; the frame number of the raw binary frame implicitly encodes the *y* coordinate of the events in the output image; the *j* coordinate in the frame has no significance in our optical setup.

Thus batches of *n* event matrices can be summed to generate one row of pixels in the output image *I*, where *n* is chosen based on the beam scan velocity and the desired pixel aspect ratio for the output image (usually 1:1):

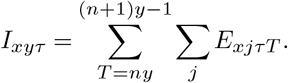

Repeating this process for all raw single-photon frames yields a complete time-of-arrival histogram image for the sample (Figure 1b).

Because our light-sheet is formed by scanning the pencil beam with a symmetric saw-tooth trajectory, each sample region is in fact imaged twice within a full beam scan (once on the up-sweep and once on the down-sweep). This means that the image formed by this process is mirrored vertically about the row *y*_*mirror*_, indicated by the blue dashed line in Figure 1b. Therefore, as a final step (“de-mirroring ”), we fold the image about *y*_*mirror*_, effectively summing the histograms for both repeated copies of each pixel. This operation can be expressed as the mapping:

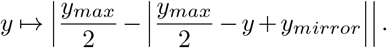

Since we were unable to synchronize our galvo sweep to the timebase of the SPAD array, *y*_*mirror*_ is not known *a priori* and must be identified through post-processing. *y*_*mirror*_ is chosen to yield the strongest correlation between the two halves that are combined in the de-mirroring operation.

Each pixel of our time-of-arrival histogram is analysed to determine its fluorescence lifetime. For this proof of concept we used the FlimJ plugin (13) to perform a pixelwise tail-fitting procedure subject to a Poisson noise model, modelling photon time-of-arrival as *N*_0_*e*^*−λt*^, where *t* is the evolved time; *H*(*t*) is the recorded histogram of photon arrival times for a given pixel; *N*_0_ is the initial number of recorded photons and *λ* is the decay constant. We note that our data acquisition and processing pipeline is equally compatible with alternative lifetime analysis methods (14, 15).

We first demonstrate the ability of our system to rapidly and simultaneously acquire fluorescence lifetime and intensity images on our DSLM-FLIM system. We imaged 300 um thick mouse lung tissue sections (fixed using 4% PFA, stained with Cyanine 3 fluorescent dye targeting the CD31 protein, and optically cleared using Ce3D clearing solution post-fixation). The tissue was injected with a solution of fluorescent beads (Bangs labs Envy Green, *ø* = 1*µ*m) to simulate the presence of foreign bodies with a similar fluorescence emission spectrum to the host tissue. The excitation laser was a single-photon 532 nm source generating ∼ 90 ps pulses at a pulse repetition rate of 80 MHz (Horiba DeltaDiode Solas-532L). The DSLM beam was swept through a *y* scan range of 290 µm at a frequency of 0.2 Hz. Figure 2a displays the reconstructed fluorescence intensity image; Figure 2b displays the same data subject to a small gamma correction to assist in visualising the full dynamic range of the data (using a mapping 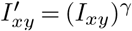, where *I*_*xy*_ is the measured intensity value at pixel (*x, y*) and here *γ* = 0.7). Figure 2c displays the corresponding fluorescence lifetime image, where a lifetime value was calculated for every pixel above a minimum threshold of 1000 photons. The green arrows indicate regions in the sample where it would be especially unclear from the intensity image alone whether the structure is tissue or a foreign body; the contrast in the lifetime image enables us to clearly discriminate between the two types of structure.

**Fig. 2.**
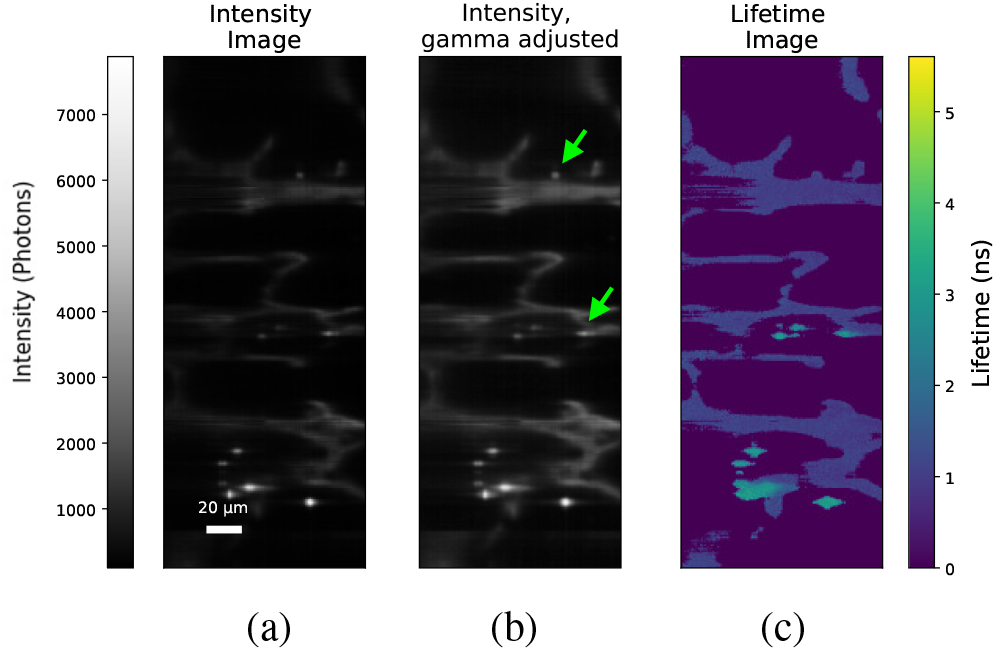
Intensity and lifetime imaging of a sample of clarified mouse lung tissue stained with Cy3 targeting the CD31 protein. The sample was also injected with a solution of fluorescent microspheres with peak emission at ∼570 nm. Left: intensity image, where the total recorded photons are summed into a single 2D image. Centre: Intensity image after modest gamma correction to better display full dynamic range. Right: Fluorescence lifetime image of the same sample. Green arrows indicate regions where the structure is ambiguous in the intensity image, but the contrast (or lack of) in the lifetime image distinguishes which structural component the feature represents. Scale-bar 20 µm.

We next demonstrated two-photon volumetric DSLM-FLIM imaging of a mixed sample of three different types of fluorescent microspheres with contrasting fluorescence life-times (Invitrogen 6 *µ*m, product code C16509; PolyAn 6.5 *µ*m, product code 11010006; PolyAn 6.5*µ*m, product code 11020006) embedded in agarose gel. Two-photon excitation was generated from a modelocked ultrashort pulse laser producing 120 fs pulses at a centre wavelength of 1040 nm (Chromacity 1040), operating at a pulse repetition rate of 80 MHz. The beam-shaping optics were designed to deliver a pencil beam with a waist diameter of 3.5 µm, and the beam was scanned across a vertical range of 370 µm. A *z*-stack was acquired by translating the sample along the optical axis of the imaging objective with a motorised stage, and combining multiple 2D+lifetime acquisitions into a 3D+lifetime stack. Figure 3 shows an individual plane and maximum intensity projection of the acquired 113 µm × 370 µm × 153 µm (*xyz*) volume, displayed using a custom colourmap indicating the lifetime of each voxel (3D rendered flyaround available as Supplementary Video 1).

**Fig. 3.**
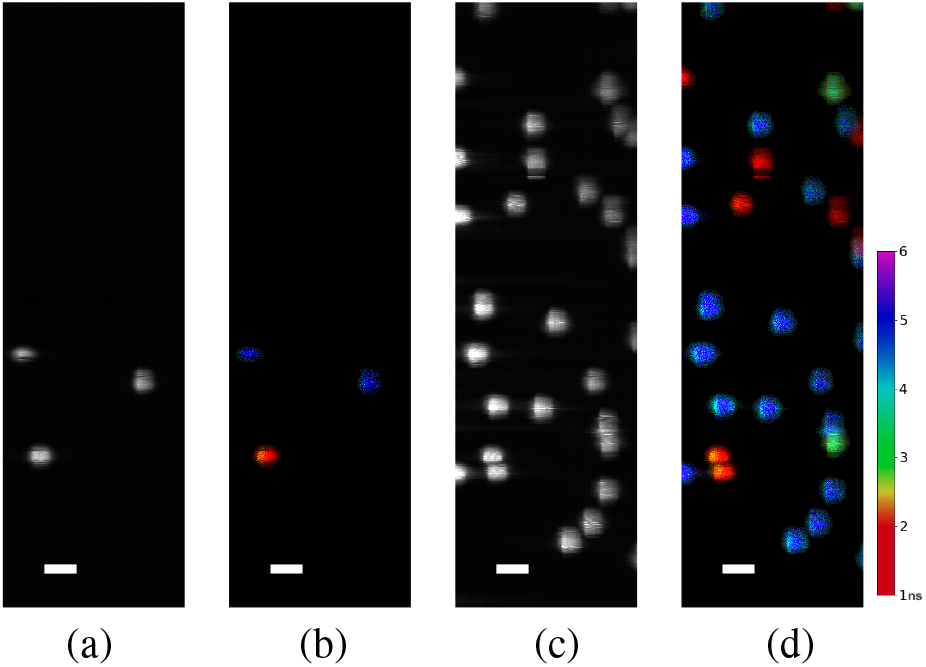
Two-photon intensity (a,c) and fluorescence lifetime (b,d) images of a single plane (a-b) and a maximum intensity projection of a 3D stack (c-d) of mixed fluorescent microspheres. Due to their very similar radii and emission spectra, the microspheres cannot be distinguished from an intensity-only image, but they can be clearly distinguished by their fluorescence lifetimes. Supplementary Video 1 shows a rendered flyaround of this dataset. Scale-bars 20 µm.

Our DSLM-FLIM results demonstrate the feasibility of our astigmatic imaging system for fast light-sheet fluorescence lifetime imaging, by enabling efficient use of the high frame-rates offered by a SPAD array camera. This is particularly crucial for two-photon FLIM, since scanless widefield two-photon excitation is prohibitively inefficient outside a few very specific scenarios (16) and hence a scan-based approach is near-essential.

Our astigmatic optical design is not directly image-forming in the *y* direction; the *y* resolution is dictated by the vertical extent of the pencil excitation beam. A narrow beam waist, therefore, yields better axial and vertical image resolution. However, the limited Rayleigh range of a propagating Gaussian beam (17) means that a narrow waist can only be maintained over a restricted lateral field of view. In all light sheet systems this trade-off constrains the *x* field of view for effective optical sectioning, but we note that in our approach the *y* resolution of the imaging system is also affected.

On the other hand, a benefit of our optical approach is that it expands the *y* field of view, and the number of effective sampled pixels. Instead of being restricted to the pixel count of the SPAD array, the achievable field of view in *y* is determined by the field of view of the objective lens itself, and the scan range of the excitation laser beam. The number of pixels sampled vertically can be selected computationally, post-acquisition, based on a trade-off between scan speed, emitted photon flux, and the required per-pixel photon budget for FLIM analysis. The vertical *resolution* is determined by the excitation beam parameters, as noted above.

Our optical approach is crucial for efficient two-photon FLIM using a SPAD array, due to our ability to accelerate acquisition times and photon-efficiency by up to two orders of magnitude. Our approach is also significant in the context of one-photon FLIM. In addition to increasing the field-of-view in *y*, the DSLM/SPAD combination offers improved signal-to-noise ratio, and the ability to implement background rejection using confocal slit detection.

We have demonstrated a proof of concept of a new astigmatic imaging approach allowing optimal co-deployment of TCSPC SPAD cameras and DSLM illumination, with either one-or two-photon excitation. We anticipate that our approach will open up a new application space of high-speed fluorescence lifetime imaging using TCSPC SPAD cameras.

## Acknowledgements

The Flimera and DeltaDiode were loaned by HORIBA Jobin Yvon IBH Ltd, and the Chromacity 1040 ultrafast laser source by Chromacity Limited, as in-kind support towards the project.

